# Highly efficient and specific genome editing in human cells with paired CRISPR-Cas9 nickase ribonucleoproteins

**DOI:** 10.1101/766493

**Authors:** Jacob Lamberth, Laura Daley, Pachai Natarajan, Stanislav Khoruzhenko, Nurit Becker, Daniel Taglicht, Gregory D. Davis, Qingzhou Ji

## Abstract

CRISPR technology has opened up many diverse genome editing possibilities in human somatic cells, but has been limited in the therapeutic realm by both potential off-target effects and low genome modification efficiencies. Recent advancements to combat these limitations include delivering Cas9 nucleases directly to cells as highly purified ribonucleoproteins (RNPs) instead of the conventional plasmid DNA and RNA-based approaches. Here, we extend RNP-based delivery in cell culture to a less characterized CRISPR format which implements paired Cas9 nickases. The use of paired nickase Cas9 RNP system, combined with a GMP-compliant non-viral delivery technology, enables editing in human cells with high specificity and high efficiency, a development that opens up the technology for further exploration into a more therapeutic role.

## Introduction

Genome modification efficiency has increasingly become the focus for new technology development as genome editing has gained more widespread adoption. CRISPR systems, naturally present in various bacterial and archeal species as adaptable immune systems, are currently being adapted for genome editing in a wide variety of cellular and multi-cellular organisms^1,2^. The system works using CRISPR-associated (Cas) nucleases that are guided to a specific DNA sequence by a complementary guide RNA, where a genome cleavage event occurs^1,2^. The primary advantage of CRISPR systems over zinc-finger nucleases (ZFNs) and transcription activator-like effector nucleases (TALENS) relates to its RNA-guided nature that enables easy programming to nearly any genomic target sequence. Designing a CRISPR guide RNA is much more affordable and practical than engineering a new ZFN or TALEN protein, leading to CRISPR’s increasing adoption as a technology^1,2^. Since its initial discovery in bacteria, CRISPR has been adapted to work in a multitude of different cell types, including yeast, plants, and mammals^2–5^.

The development of CRISPR systems has not stopped at just expanding applications to new varieties of cell types and model organisms. As therapeutic applications have developed, concerns have been raised about the specificity of CRISPR-based genome modifications. With enough complementarity between the intended site and a genomic site at another location, off-target cleavage has been shown to occur with the most popular variety of Cas nuclease from *Streptococcus pyogenes* (SpCas9)^6–8^. This off-target activity has prompted the development of multiple modifications to make high efficiency versions of SpCas9, which possess a relaxed binding efficiency shown to result in higher on-target fidelity without loss in overall cleavage efficiency^9,10^. Another solution that addresses the off-target problem of SpCas9 is the development of paired SpCas9 nickases. In the paired nickase approach, SpCas9 has been modified to render one of its nuclease domains non-functional, causing the protein to make a single-stranded nick rather than a double stranded break (DSB). Paired with a second Cas9 nickase targeting a proximal location, these two single-stranded nicks cause a DSB to occur^11,12^. Off-target sites modified using both paired gRNA are exceedingly rare, and any off-target sites modified by a single Cas9 nickase are accurately repaired by the cell to render off-target mutagenesis undectable.^6,11–14^

The delivery of guide RNA and Cas9 protein to cells is another aspect that has to be considered when considering the efficiency of CRISPR systems. Conventionally, plasmids that encode the Cas9 protein and guide sequences have been transfected into cells via either lipofection or electroporation. These methodologies have the drawbacks of random integration of the plasmids into the host genome and prolonged expression of the nucleases and guide sequences, which can cause immune responses and greatly increase off-target effects, respectively, limiting the effectiveness of the technology for therapeutic purposes^15^. In order to improve upon plasmid-based limitations, complexes comprised of purified recombinant Cas9 protein and guide RNA (RNP) have been developed for direct delivery to a multitude of different human cell types. These RNPs have been shown to cut genomic DNA rapidly after delivery and degrade rapidly, thereby decreasing off-target concerns associated with the stronger Cas9-gRNA expression characteristic of plasmid-based systems^15–19^.

In this study, we demonstrate the effective combination of paired nickase and RNP formats in genome editing of human cells, including primary T cells, with high efficiency and specificity.

## Materials and Methods

### Cas9 Proteins, tracrRNA, crRNA and sgRNA

Recombinant SpCas9-D10A nickase proteins were expressed and purified from *E. coli*, and now are available in MilliporeSigma. Wild-type SpCas9 protein, tracrRNA, crRNA, and single-gRNA (sgRNA) were all sourced from MilliporeSigma.

### RNP Formation

To form the RNPs, first all three parts (Cas9 Protein, tracrRNA and crRNA) or two parts (Cas9 protein and sgRNA) were resuspended to a concentration of 30μM in either the supplied resuspension solution or 10 mM Tris buffer with a pH of 7.5. They were then assembled in an 11 μL mixture at a molar ratio of 5:5:1 (crRNA:tracrRNA:Cas9 protein) or a 4 μL mixture at 3:1 (sgRNA:Cas9 protein) and left at room temperature for 5-15 minutes immediately before use.

### In vitro DNA cleavage

*In vitro* activity of Cas9 nickase protein was evaluated by measuring the relaxation of supercoiled pUC19 plasmid DNA (MilliporeSigma). The reaction (30 μL) contained 3nM DNA and Cas9-nickase RNP in 20 mM HEPES, 100 mM NaCl, 5 mM MgCl_2_, 0.1 mM EDTA, pH 6.5, for 30 min at 37°C. Samples were electrophoresed by running them on a 1% agarose gel (MilliporeSigma) in TAE buffer (MilliporeSigma). Percent digestion was calculated by measuring the intensity of supercoiled and relaxed bands with ImageQuant LAS 400 (GE), referred to their percentage in the same assay without gRNA, the negative control.

### Cell Culture

The human K562 cells were acquired from ATCC and were maintained in a modified DME (MilliporeSigma) with 10% FBS (MilliporeSigma), 2% Glutamine (MilliporeSigma), and an antibiotic mix of penicillin and streptomycin (MilliporeSigma). The K562 cells were split 1 day prior to transfecting and transfections were performed at ~1 million cells per ml.

Human primary CD8+ T cells were purchased from STEMCELL Technologies Inc. Cells were maintained in Stemline® T Cell Expansion Medium (MilliporeSigma) supplemented with 10% human AB serum (MilliporeSigma), 1x GlutaMAX™ (Gibco), 8ng/mL IL-2 (Gibco), and 50 μM β-mercaptoethanol (MilliporeSigma). Cells were stimulated with Dynabeads™ Human T-Expander CD3/CD28 (Gibco) 7 days prior to nucleofection. Cells were cultured in the presence of Dynabeads™ post nucleofection according to manufacturer’s protocol.

Human peripheral blood mononuclear cells (PBMCs) were isolated from leukapheresis blood product (Hemacare), then suspended in X Vivo 15 medium (Lonza) with 100U/ml of IL-2 (R&D Systems). T Cells were stimulated with Dynabeads™ Human T-Expander CD3/CD28 (Gibco). The cells were cultured in GRex (Wilson Wolfe) 6 well dishes at 4-5 million per ml for 2 to 3 days before transfection with the MaxCyte^®^ Flow Electroporation™ system.

### Transfection

For K562 cells, enough cells were obtained for approximately 350K cells per transfection, while for the CD8+ human primary T cells 500K cells per transfection were used. Transfection was done using the 4D-Nucleofector™ System (Lonza) with the entire RNP mix added to 100μl of cells before using the manufacturer recommended protocol for the cell line. For the paired nickase tests, two nickase RNPs (each contains one of the paired crRNAs) were formed separately and both added to the cells simultaneously immediately before transfection. The cells were plated on 12-well plates with 1.5ml complete growth media per well and incubated at 37°C before DNA analysis. Transfection of PBMCs was completed using the MaxCyte electroporation system.

### CEL-I Analysis of CRISPR/Cas9 Cleavage Activity

Genomic DNA was extracted from the cells using QuickExtract DNA Extraction Solution (Epicentre) following the protocol of the manufacturer and the target sites were PCR amplified using the primers listed in the supplement ordered from MilliporeSigma. The CEL-I Assay, a mismatch-nuclease based assay that detects single-base mismatches or small insertions or deletions, was performed using the Surveyor Mutation Detection Kit (IDT) according to its instructions. First, the PCR amplicons go through a denaturing and annealing step in the thermocycler after amplification to form a heteroduplex, followed by a digestion with the Nuclease and Enhancer proteins at 42°C before being electrophoresed on a 10% TBE Gel (Thermofisher). The gel was then stained in 100 ml 1x TBE buffer with 2 μl of 10 mg/ml ethidium bromide for 5 min, then washed with 1x TBE buffer and visualized with a UV illuminator. The resulting bands were analyzed using Image J software.

### TOPO Cloning

The PCR amplicon obtained for the CEL-I assay was also ligated with the pCR™2.1-TOPO® vector (Thermofisher) using the manufacturer’s recommended protocol. The ligation products were then used for transformation into One Shot™ TOP10F’ Chemically Competent *E. coli* (Thermofisher) and 20 colonies were sequenced via Sanger sequencing using universal primer M13R.

### Next-Generation Sequencing (NGS)

JumpStart™ REDTaq® ReadyMix™ Reaction Mix (MilliporeSigma) along with primers flanking the genomic cut site of EMX1 and PD1 were used for PCR amplification. Primers were tagged with a partial Illumina adapter sequences using the primers found in the supplement. The thermal cycling conditions included a heat denaturing step at 95°C for 5 minutes followed by 34 cycles of 95°C for 30 s, anneal at 67.7°C for 30 s, and extension at 70°C for 30 s. Amplification was followed by a final extension at 70C for 10 min and a cool down to 4°C.

A limited-cycle PCR was carried out to index the amplified PCR product. A total reaction volume of 50 μL included 25 μL JumpStart™ REDTaq® ReadyMix™ Reaction Mix (MilliporeSigma), 5 μL of amplified PCR product, 10 μL H2O, and 5 μL each of 5 μM Nextera XT Index 1 (i7) and Index 2 (i5) oligos. The thermal cycling conditions consisted of an initial heat denature at 95°C for 3 min, followed by 8 cycles of 95°C for 30 s, 55°C for 30 s, and 72°C for 30 s. A final extension was carried out at 72°C for 5 min and the reaction was cooled down to 4°C. PCR purification was carried out using AxyPrep™ Mag PCR beads (Corning) and 25 μL of indexed sample at an 8:1 bead-to PCR ratio. DNA was eluted in 25 μL of 10mM Tris.

PicoGreen fluorescent dye (Invitrogen) was used for quantification of indexed samples. Purified indexed PCR was diluted to 1:100 with 1xTE. PicoGreen was diluted according to manufacturer’s protocol (50 μL PicoGreen + 10mL 1xTE). Equal volume of diluted PicoGreen was added to the diluted indexed PCR sample yielding a final 1:1 dilution ratio in a fluorescence plate reader. Samples were excited at 475 nm and read at 530 nm. All samples were normalized to 4nM with 1xTE, and 6 μL of each normalized sample was collected and pooled.

Stock 10M NaOH was serial diluted with H_2_O to yield a final 1:100 dilution (.1M NaOH) the day of library preparation. To denature the DNA, 5 μL of .1M NaOH and 5μL of the pooled 4nM library were mixed together in an Eppendorf tube and incubated at RT for 5 minutes. 990 μL of cold Illumina HT1 buffer was added to the denatured DNA, yielding a 20pM pooled, denatured library. PhiX (20pM) was thawed and 30 μL was transferred to a fresh tube. 570 μL of the 20pM library was added to the PhiX resulting in 5% PhiX for library diversification, quality control for cluster generation, sequencing, and alignment. This was mixed and heat shocked at 96C for 2 minutes and then immediately placed on ice. A 300 cycle v2 Miseq reagent cartridge was thawed in an ice water bath and inverted to mix. A Miseq v2 flow cell was rinsed thoroughly with water and ethanol and dried with a Kimwipe making sure that all liquid was removed and no salt was detected on the surface of the cell. A p1200 tip was used to pierce the foil of well 17 of the thawed reagent cartridge. Using a new tip, 600 μL of the PhiX containing library was added to well 17. Following the run, .bam files were used for analysis with IGV software.

### Sanger Trace Analysis to Quantify Insertions / Deletions (indels)

Genomic DNA was extracted from the cells using QuickExtract DNA Extraction Solution (Epicentre) or PureLink Solution (Invitrogen) following the protocol of the manufacturer. Target-specific PCR products were sequenced by GENEWIZ and Sanger traces analyzed for increased sequence heterogeneity to characterize indel levels and location.

### Flow Cytometry

About 1 million cells were washed once with flow cytometry staining buffer (BD Biosciences) by centrifugation at 500 g for 5 min, resuspended in 100 μL of the same buffer, and then incubated for 30 minutes with 20 μL of anti-PD1 antibody (BD Biosciences). The cells were washed once again. After incubation with 7-AAD viability dye (eBioscience) for 5 min, samples were run on Beckman Coulter CytoFLEX flow cytometer. Analysis of the flow cytometry data was done by using FlowJo software, version 10.5.3

## Results

### Genome editing with paired SpCas9-nickase RNPs in human K562 cells

Our first step in evaluating the paired nickase RNP format was to determine if it was able to cleave DNA *in vitro*. We found that when the SpCas9 nickase was formed into an RNP it was able to relax on average 90% of supercoiled pUC19 plasmid DNA that it targeted, compared to the immeasurable activity of the individual components alone (Supplementary Figure S1).

Next we tested paired nickase RNPs in human K562 cells, known in our hands to be an excellent first pass option for evaluating new ZFN and CRISPR molecular systems. We first tested multiple crRNA targets for the EMX1 locus using SpCas9 RNPs to determine if targeting these sites with a wild-type SpCas9 protein was able to generate any genomic insertion / deletions (indels), and found multiple viable target sites to move forward with (data not shown). We then delivered the paired SpCas9-nickase RNP complexes via nucleofection into K562 cells and harvested the genomic DNAs for detection of genomic modification.

TOPO cloning assays detected sequence modification at the expected target sites associated with the paired nickase RNP treatments (data not shown). Next a CEL-I assay was performed, and detected efficient genome editing with paired nickase RNPs, at levels similar to those from wide-type SpCas9 RNP; as expected, no indels were detected from nickase RNPs with targeted using a single crRNA (Figure 1A).

**Figure 1.**
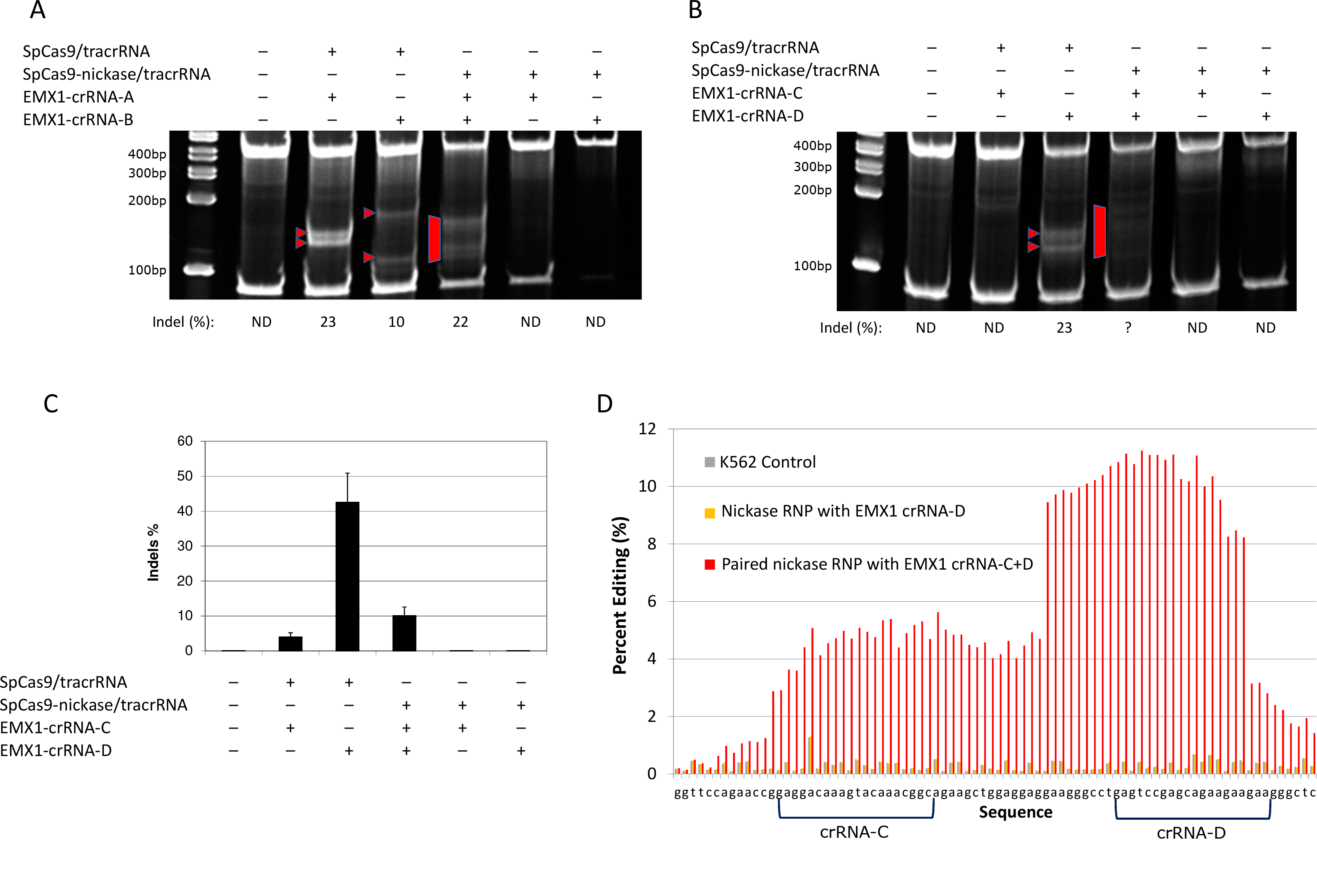
Genome editing with paired SpCas9 Nickase RNP in K562 cells. Paired SpCas9 nickase RNPs, along with SpCas9 RNPs, were transfected into K562 cells. (A) CEL-I analysis for RNP mediated indels at the EMX1 Locus. The cleaved fragments from SpCas9 RNP treated samples were indicated by red triangles, and the cleaved fragments from paired SpCas9-nicakse RNP treated samples were indicated by red trapezoid. Data are representative of three biological replicates. ND, not detected. (B) CEL-I analysis for RNP mediated indels at the other EMX1 Locus. (C) NGS analysis for RNP mediated indels at the same Locus as B. Averages from three biological replicates are plotted with error bars representing one standard deviation. (D) The percent editing of each single nucleotide around paired crRNAs targeted region measured by NGS analysis. Data are representative of three biological replicates.

We evaluated multiple loci using paired nickase RNPs, and for some samples the CEL-I assay was not able to yield clear results due to the smeared bands in the gel (one example is shown in Figure 1B (lane 5). We hypothesized that indels caused by the paired nickase RNPs happen across a much wider region, compared to those caused by a single SpCas9 nuclease. In order to measure accurate indels resulting from paired nicks, we applied Next Generation Sequencing (NGS) which clearly detected expected genome modifications (Figure1C). NGS analysis confirmed that the paired nickase RNPs induced genome modifications in between the paired nicking sites. Moreover, Cas9-nickase RNPs guided by a single crRNA sequence yielded NGS results indistinguishable from the control DNA, confirming the necessity of two proximal nicks to generate detectable indels (Figure 1D) and suggesting low chances of genome modification at off-target sites modified by Cas9-nickases guided by a single crRNA.

### Genome editing with paired SpCas9-nickase RNPs in primary human T cells and PBMCs

Next we investigated the ability of our paired SpCas9-nickase RNP system to modify a more clinically relevant human genomic target encoding the protein programmed death-1 (PD-1). Targeting this gene has been shown to be an effective approach for checkpoint inhibition, which can have applications in developing T-cell based therapies^20^. Before testing in T cells, we again tested in human K562 cells to determine the efficiency of particular gRNA designs in the absence of potential cellular delivery limitations that might be associated with unique aspects of T-cells sourced from various patients and vendors. We designed three pairs of PD-1 crRNAs, which all yielded detectable genome modification at the target loci when complexed and delivered with SpCas9 nickases and tracrRNAs (data not shown). When applying these three paired nickase RNPs into human primary CD8+ T cells, we observed either undetectable or very weak genome editing at targeted loci. We then optimized our nickase RNP system with chemically modified single gRNA (mod-sgRNA), instead of two part gRNA (tracrRNA + crRNA). Mod-sgRNA worked significantly better than two parts tracrRNA + crRNA for paired nickase RNP in human primary CD8+ T cells, measured by Sanger trace analysis (Supplementary Figure S2). Mod-sgRNAs were then used for all following experiments in T cells.

To demonstrate feasibility of using nickase RNPs for therapeutic applications, we transfected the reagents into human primary hematopoietic cells using a clinically validated, GMP-compliant electroporation technology: Paired nickase RNPs were delivered into human primary PBMCs using the MaxCyte GTx Flow Electroporation system. About 50% indels were generated by nickase RNP pair 2 mod-sgRNAs (PD-1 mod-sgRNA-C+D), even higher than those from the best SpCas9-RNP (with PD-1 mod-sgRNA-C) (Figure 2A). Similar to previous results, no indels were detected from either nickase RNP targeted by a single mod-sgRNA, while high indels were found in the paired delivery (Figure 2B).

**Figure 2.**
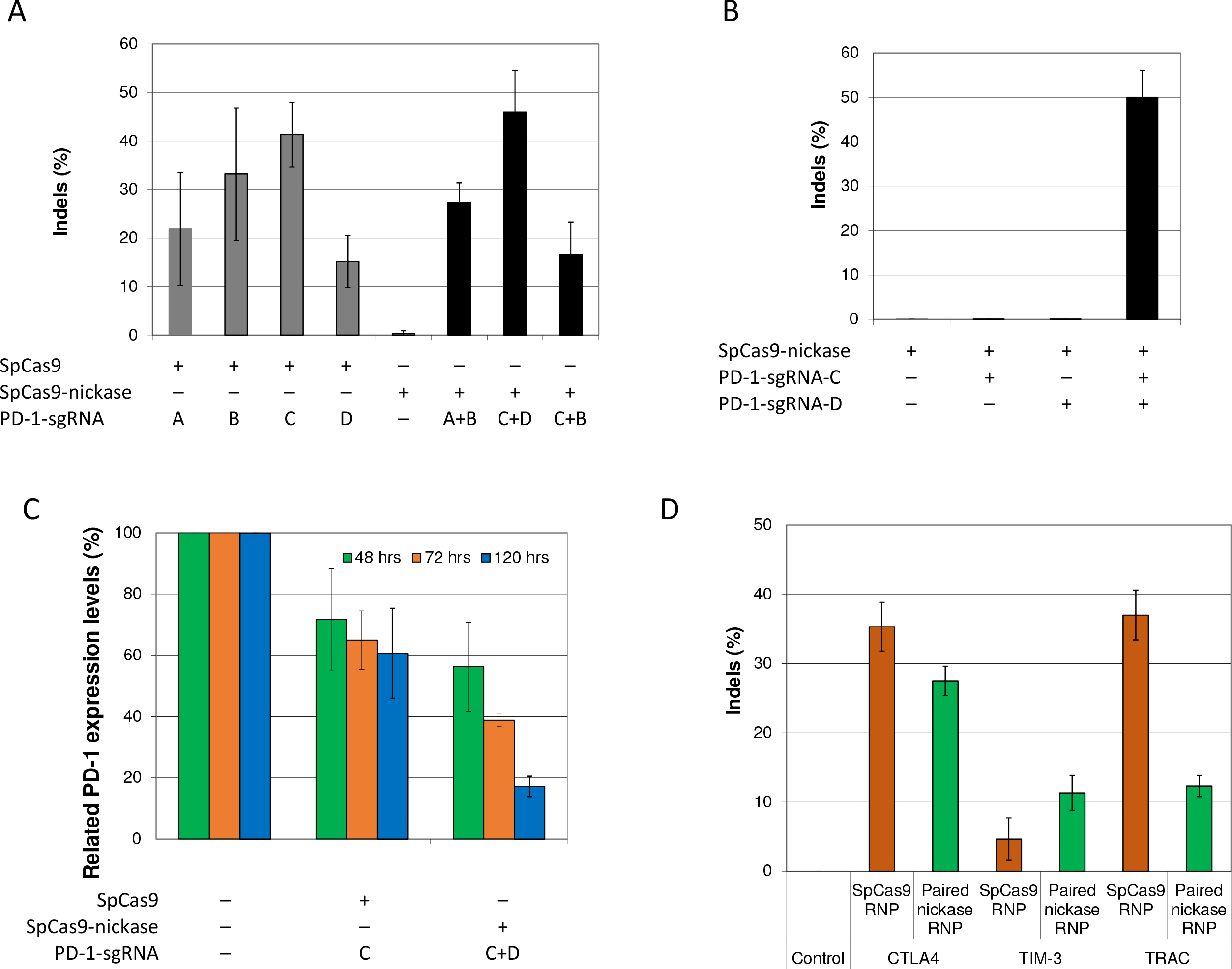
Genome editing with paired nickase RNP in human primary T cells and PBMCs. (A) Paired SpCas9 nickase RNP on variant PD-1 loci, along with SpCas9 RNPs, were delivered into human primary PBMCs through MaxCyte electroporation. Pair 1: PD-1 mod-sgRNA-A+B; pair 2: PD-1 mod-sgRNA-C+D; pair 3: PD-1 mod-sgRNA-C+B. Genomic DNAs were harvested 72 hours post-electroporation for indels analysis. Averages from three biological replicates are plotted with error bars representing one standard deviation. (B) Paired SpCas9 nickase RNP, along with nickase RNP with single sgRNA, were delivered into Human primary T cells. Genomic DNAs were harvested 72 hours post-electroporation for indels analysis. Averages from three biological replicates are plotted with error bars representing one standard deviation. (C) Paired nickase RNP, along with nickase RNP containing single sgRNA or SpCas9 RNP, were delivered into human primary PBMCs. At variant time points, cells were stained with anti-PD-1 antibody for flow cytometry analysis. Averages from three biological replicates are plotted with error bars representing one standard deviation. (D) Paired SpCas9 nickase RNP, along with SpCas9 RNP, on various targets (Paired nickase RNP with CTLA4 pair 2, SpCas9 RNP with CTLA4 sgRNA-D, paired nickase RNP with TIM-3 pair 3, SpCas9 RNP with TIM-3 sgRNA-F, paired nickase RNP with TRAC pair 2, SpCas9 RNP with TRAC sgRNA-D) were delivered into Human primary CD8+ T cells. Genomic DNAs were harvested 120 hours post-electroporation for indels analysis. Averages from three biological replicates are plotted with error bars representing one standard deviation.

Next, we determined if genome modifications to the PD-1 locus by paired nickase RNPs are associated with reductions in PD-1 protein expression levels in cells. Paired nickase RNP (with PD-1 mod-sgRNA pair 2) was again delivered to human primary PBMCs through MaxCyte electroporation. At variant time points, cells were stained with anti-PD-1 antibody for flow cytometry analysis. As shown in Figure 2C (and Supplementary Figure S3), at 48 hours after paired nickase RNP treatment, a reduction of PD-1 protein expression levels was easily detected; and further reduction occurred with longer time treatment. Less than 20% PD-1 FACS signal was left 120 hours following electroporation of paired nickase RNPs. Unexpectedly, paired nickase RNPs were more effective than SpCas9-RNP at reducing PD-1 protein FACS signal, especially after 120 hours treatment. Our previous data indicated indels caused by paired nickase RNPs happens in a wide region spanning both guide targets. Compared with typical SpCas9 nucleases using one gRNA, this wider region of indels caused by paired nickase RNPs could lead to more efficient knockout of gene functionality by deletion of additional amino acid residues or changes in mRNA structure which enhance mRNA destruction mechanisms such as non-sense mediated decay.

In addition to PD-1, cytotoxic T-lymphocyte protein 4 (CTLA4), T-cell immunoglobulin and mucin-domain containing-3 (TIM-3; also called hepatitis A virus cellular receptor 2, HAVCR2) and T-cell receptor alpha constant (TRAC) are all emerging targets in the cancer immunotherapy landscape^21–24^. Sets of paired gRNAs were designed for these targets and pre-screened in K562 cells (data not shown). The best of these gRNA pairs generated efficient genome editing on all three gene targets in human primary CD8+ T cells (Figure 2D).

## Discussion

The CRISPR-Cas9 system is still in the early stages of development for clinical applications. The full potential of the technology is still being developed with different versions and methods coming into relevance as researchers across the world explore variations. The paired nickase version of Cas9 is one such variation that uses a modified Cas9 to make two single-stranded complementary cuts in DNA to form a DSB. Because this method uses two independent guides, the production of permanent genome modifications is highly site specific^11,12^. Here we report the successful combination of paired Cas9 nickases and RNP-based CRISPR delivery^15,17^. We have shown above that our paired nickase RNP system has high activity at a variety of targets in multiple cell types. In addition, our paired nickase RNP system also only produces genomic indels when Cas9 nickase RNPs are paired and co-delivered into the cell with two proximal gRNA, eliminating the chance of off-target indel activity. The combination of a highly specific and highly efficient Cas9 system can allow the technology to be better suited for therapeutic or clinical use. A previous publication (Gopalappa et. al., 2018) also suggested that paired Cas9-D10A nickases sometimes could be more efficient than individual nuclease for gene disruption, although their results were based on a plasmid Cas9-nickase system rather than our RNP system^14^.

One potential application for paired nickase RNPs is to modify gene targets to improve *ex vivo* cell-based immunotherapy, a powerful treatment option that harnesses the immune system to fight cancer, infection, and other diseases. Conventional immunotherapy comprises the use of substances such as vaccines, monoclonal antibodies, cytokines, etc. to stimulate or suppress the immune system and other compounds. In recent years, genome editing is being used to modify the DNA of cells to engineer better functioning cells for use in immunotherapy^22–25^. The PD-1 receptor, one of the key checkpoint inhibitors for cancer immunotherapies, is bound on activated T-cells by the ligand PD-L1 on some tumor cells as a negative regulator^20^. The antitumor effect of tumor reactive T-cells can be improved by blocking the PD-1 receptor, increasing the immune response instead of inducing cell death directly^25^. This checkpoint inhibition has been accomplished with a variety of methods so far, ranging from blocking antibodies to knockout using zinc finger nucleases (ZFNs) or transcription activator-like effector nucleases (TALENS)^20^. Now with the development of Cas9 systems, both plasmid and RNP versions of Cas9 systems have also been demonstrated to be effective in targeting the site before^18,20^.

We have demonstrated here that a paired nickase RNP delivery system is able to produce targeted genomic modifications in multiple immune-related genes, including PD-1, CTLA4, TIM-3 and TRAC in human primary T-cells with high specificity and efficiency. Further supporting clinical feasibility, efficient gene knockout was exhibit following delivery via a GMP compliant, scalable electroporation platform. As stated above, the increased specificity and efficiency with our paired nickase RNP system, combined with optimized cellular delivery methods, create a promising alternative for research and therapeutic applications.

## Acknowledgements

We gratefully thank Tim Seebeck and Daniel Lynch for technical assistance on NGS; Patrick Sullivan and James Brady for R&D administrative support, and MilliporeSigma, a business of Merck KGaA, Darmstadt, Germany, for financial support.

## Supplementary Material

Supplementary Materials are available online

