## Supplementary Materials for "Highly efficient and specific genome editing in human cells with paired CRISPR-Cas9 nickase ribonucleoproteins"

**This file includes:**
Tables S1 to S3
Figures S1 to S3

**Table S1**. crRNA/mod-sgRNA list

| Target | crRNA/mod-sgRNA | Target sequence |
|---|---|---|
| EMX1 | EMX1-crRNA-A | TGCGCCACCGGTTGATGTGA |
| | EMX1-crRNA-B | TTGCCACGAAGCAGGCCAAT |
| | EMX1-crRNA-C | GCCGTTTGTACTTTGTCCTC |
| | EMX1-crRNA-D | GAGTCCGAGCAGAAGAAGAA |
| | EMX1-crRNA-E | CACGAAGCAGGCCAATGGGG |
| | EMX1-crRNA-F | GGGGCACAGATGAGAAACTC |
| | EMX1-crRNA-G | TGAAGGTGTGGTTCCAGAAC |
| | EMX1-crRNA-H | TCACCTGGGCCAGGGAGGGA |
| | EMX1-crRNA-I | AGGTGTGGTTCCAGAACCGG |
| | EMX1-crRNA-J | CAAACGGCAGAAGCTGGAGG |
| PD-1 | PD-1-crRNA-A / PD-1-sgRNA-A | GCGTGACTTCCACATGAGCG |
| | PD-1-crRNA-B / PD-1-sgRNA-B | GCAGTTGTGTGACACGGAAG |
| | PD-1-crRNA-C / PD-1-sgRNA-C | GACAGCGGCACCTACCTCTG |
| | PD-1-crRNA-D / PD-1-sgRNA-D | GGGCCCTGACCACGCTCATG |
| CTLA4 | CTLA4-sgRNA-A | TTTGAACCCACACAGAATCA |
| | CTLA4-sgRNA-B | CCTTGGATTTCAGCGGCACA |
| | CTLA4-sgRNA-C | GGAGCGGTGTTCAGGTCTTC |
| | CTLA4-sgRNA-D | GCACAAGGCTCAGCTGAACC |
| | CTLA4-sgRNA-E | CCTTGTGCCGCTGAAATCCA |
| | CTLA4-sgRNA-F | GCAAAGGTGAGTGAGACTTT |
| TIM-3 | TIM-3-sgRNA-A | GGCGGCTGGGGTGTAGAAGC |
| | TIM-3-sgRNA-B | TGGTGCTCAGGACTGATGAA |
| | TIM-3-sgRNA-C | TGCCCCAGCAGACGGGCACG |
| | TIM-3-sgRNA-D | TGGTGCTCAGGACTGATGAA |
| | TIM-3-sgRNA-E | ACGTTGCCACATTCAAACAC |
| | TIM-3-sgRNA-F | CTAAATGGGGATTTCCGCAA |
| TRAC | TRAC-sgRNA-A | CAGGGTTCTGGATATCTGT |
| | TRAC-sgRNA-B | AACAAATGTGTCACAAAGTA |
| | TRAC-sgRNA-C | AGAGTCTCTCAGCTGGTACA |
| | TRAC-sgRNA-D | ACAAAACTGTGCTAGACATG |
| | TRAC-sgRNA-E | GAGAATCAAAATCGGTGAAT |
| | TRAC-sgRNA-F | CTTCAAGAGCAACAGTGCTG |

**Table S2**. crRNA/mod-sgRNA pair designs for paired nickase RNP

| Target | Pair | crRNA/mod-sgRNA |
|---|---|---|
| EMX1 | Pair 1 | EMX1-crRNA-A + B |
| | Pair 2 | EMX1-crRNA-C + D |
| | Pair 3 | EMX1-crRNA-A + E |
| | Pair 4 | EMX1-crRNA-F + G |
| PD-1 | Pair 1 | PD-1-crRNA-A + B / PD-1-sgRNA-A + B |
| | Pair 2 | PD-1-crRNA-C + D / PD-1-sgRNA-C + D |
| | Pair 3 | PD-1-crRNA-C + B / PD-1-sgRNA-C + B |
| CTLA4 | Pair 1 | CTLA4-sgRNA-A + B |
| | Pair 2 | CTLA4-sgRNA-C + D |
| | Pair 3 | CTLA4-sgRNA-E + F |
| TIM-3 | Pair 1 | TIM-3-sgRNA-A + B |
| | Pair 2 | TIM-3-sgRNA-C + D |
| | Pair 3 | TIM-3-sgRNA-E + F |
| TRAC | Pair 1 | TRAC -sgRNA-A + B |
| | Pair 2 | TRAC -sgRNA-C + D |
| | Pair 3 | TRAC-sgRNA-E + F |

**Table S3**. Primer list

| Assay | Target | Primer | Sequence |
|---|---|---|---|
| CEL-1 | EMX1 | EMX1-forward | ATGGGAGCAGCTGGTCAGAG |
| | | EMX1-reverse | CAGCCCATTGCTTGTCCCT |
| NGS | EMX1 | NickFOR-ILLUMIEMX1 | TCGTCGGCAGCGTCAGATGTGTATAAGAGACAGNNNNNNGGCCTCCTGAGTTTCTCATCTGTGC |
| | | NickREV-ILLUMIEMX1 | GTCTCGTGGGCTCGGAGATGTGTATAAGAGACAGNNNNNNNTGACTCCAGGCCTCCCCAAA |
| | PD-1 | NickFOR-ILLUMIPD1 | TCGTCGGCAGCGTCAGATGTGTATAAGAGACAGNNNNNNGGACAACGCCACCTTCACCTGC |
| | | NickREV-ILLUMIPD1 | GTCTCGTGGGCTCGGAGATGTGTATAAGAGACAGNNNNNNNCTACGACCCTGGAGCTCCTGAT |
| Sanger trace analysis | PD-1 | PD-1-forward | GGACAACGCCACCTTCACCTGC |
| | | PD-1-reverse | CTACGACCCTGGAGCTCCTGAT |
| | CTLA4 | CTLA4-forward | CCCTTGTACTCCAGGAAATTCTCCA |
| | | CTLA4-reverse | ACTTGTGAGCTCATCCTGAAACCCA |
| | TIM-3 | TIM-3-forward | TCATCCTCCAAACAGGACTGC |
| | | TIM-3-reverse | TGTCCACTCACCTGGTTTGAT |
| | TRAC | TRAC-forward | TCAGGTTTCCTTGAGTGGCAG |
| | | TRAC-reverse | TGGCAATGGATAAGGCCGAG |

### Supplementary Figure Legends

**Figure S1.** ***In vitro*** **DNA cleavage with SpCas9-nickase RNPs.** Supercoiled plasmid pUC19 was incubated with the SpCas9-Nickase RNP complex or SpCas9-nickase proteins only (negative control). Percent digestion was calculated by measuring the intensity of supercoiled and relaxed bands. ND, not detected.

**Figure S2. Comparison of editing efficiencies between paired nickse RNPs with mod-sgRNA and paired nickase RNPs with two parts tracrRNA+crRNA in human primary T cells.** Paired nickse RNPs containing either mod-sgRNA or two parts tracrRNA+crRNA were nucleofected into human primary CD8+ T cells. Genomic DNAs were harvested 72 hours post-nucleofection for indels analysis. Averages from three biological replicates are plotted with error bars representing one standard deviation.

**Figure S3. Data are representative of three biological replicates in Figure 2C.**

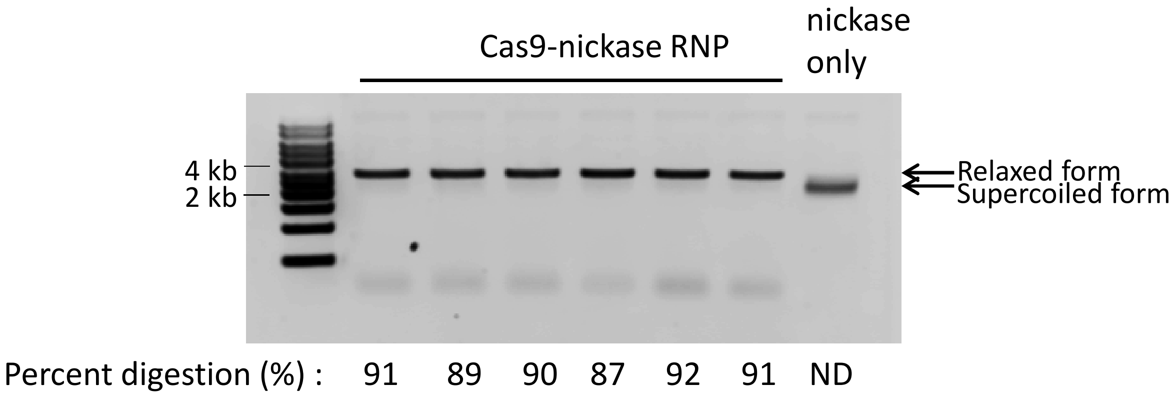


Supplementary Figure S1

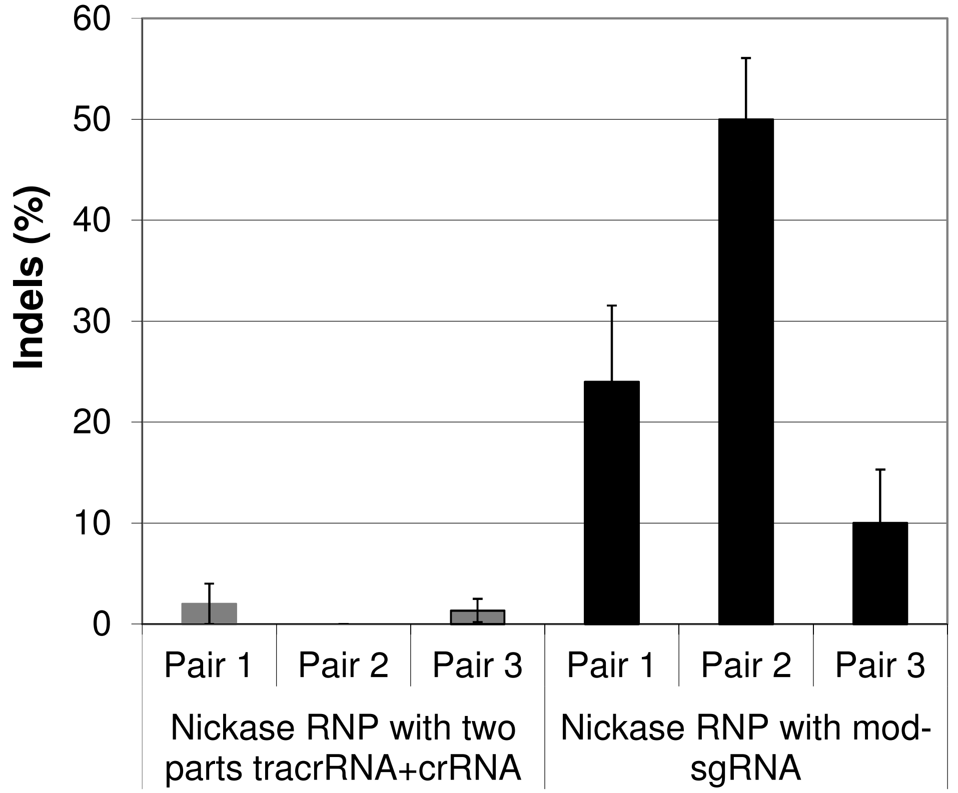


Supplementary Figure S2

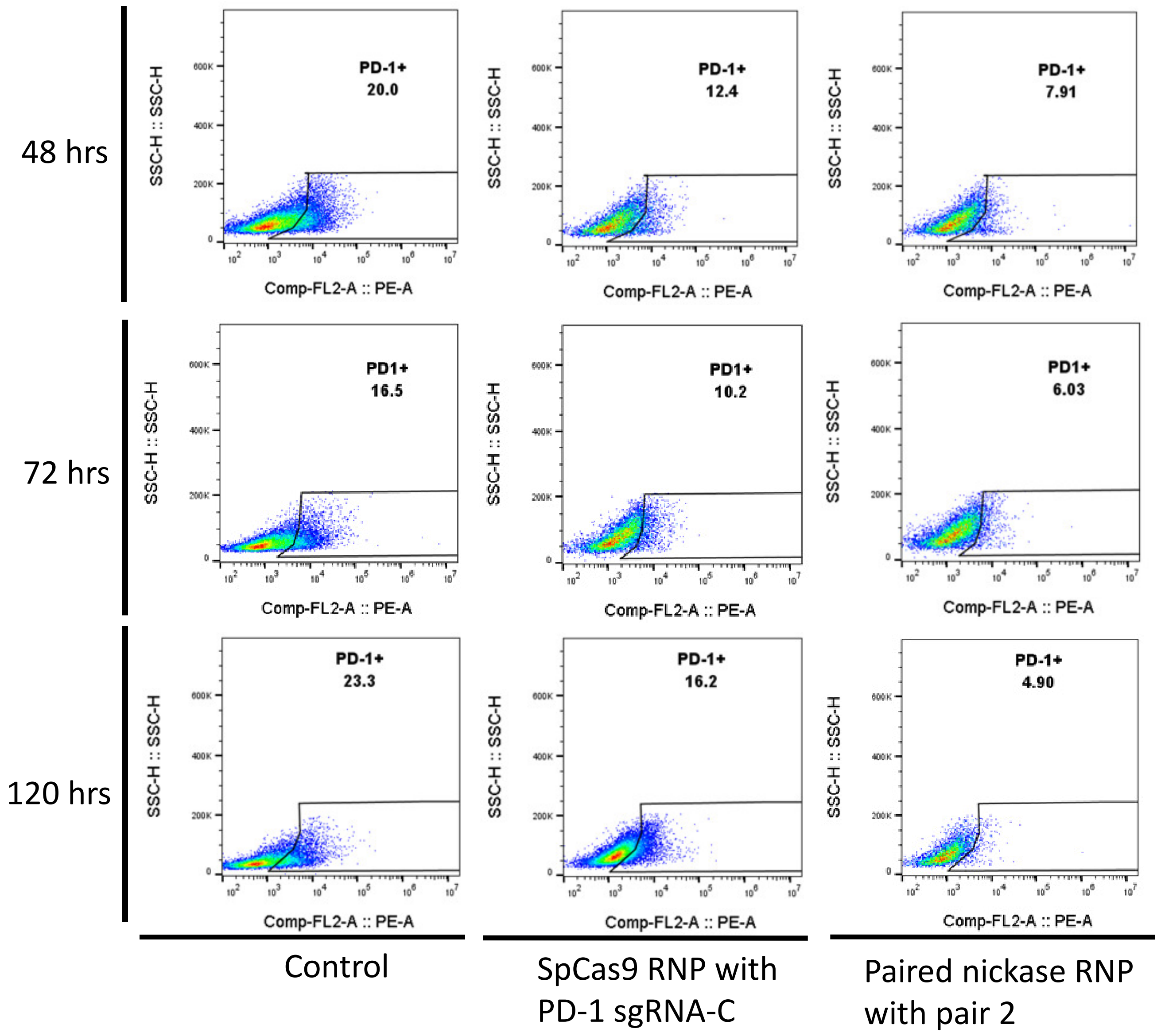


Supplementary Figure S3
